# Task-specific preferences guide forward planning in sequential decision-making

**DOI:** 10.64898/2026.09.25.754065

**Authors:** Alex Lepauvre, Florian Ott, Stefan Kiebel

**Affiliations:** Department of Psychology, Technische Universität Dresden, Dresden, Germany; Centre for Tactile Internet with Human-in-the-Loop, TU Dresden, Germany

## Abstract

Everyday problems often require choosing among actions with different future consequences, even though fully evaluating those consequences is often computationally infeasible. The human brain is remarkable in its ability to solve such complex problems efficiently. In this paper, we hypothesize that this capacity arises from a principle inspired by Bayesian accounts of perception and sensorimotor control: evidence derived from effortful forward planning is integrated with task-specific preferences akin to priors, weighted by the uncertainty of these preferences. To test this hypothesis, we used a sequential decision-making task with a relatively large state space. Model comparison showed that participants’ deviations from optimal forward planning are best captured by task-specific preferences, rather than by unsystematic errors, recent action-history effects, or offer-context biases. Moreover, the influence of forward planning on participants’ responses depended on preference uncertainty, as evidenced by both choice behavior and reaction times, and preferences were not arbitrary but instead supported above-stochastic performance even in the absence of forward planning. Together, these results are consistent with a Bayesian-inspired interpretation of decision making, in which costly forward planning is integrated with state-dependent priors or preferences. These preferences may provide an inexpensive, quickly available first-pass solution, which is complemented by forward planning when preference-based guidance is uncertain.

## 1 Introduction

A central idea in Bayesian accounts of perception and sensorimotor control is that the brain does not rely on current sensory evidence alone (Knill and Pouget 2004; Lee and Mumford 2003; Pouget et al. 2013; Clark 2013; Friston 2010) because sensory evidence is often noisy, incomplete, or costly to acquire with high precision. Prior information can reduce this uncertainty by providing an efficient summary of regularities learned from experience and from the statistical structure of the environment. Bayesian accounts provide an influential framework for describing this integration of current evidence and prior information. In this framework, the influence of prior information depends on uncertainty: when current evidence is unreliable, behavior should rely more strongly on the prior. Conversely, when current evidence is reliable, it should dominate behavior. For example, Körding and Wolpert (2004) provided a canonical demonstration of this principle, showing that participants combine learned priors with sensory feedback according to their relative uncertainty.

A similar logic may apply to decision-making based on forward planning, where choices require evaluating the future consequences of currently available actions. Here, we use the term forward planning in a computational sense: it refers to the estimation of future action consequences, not necessarily to a consciously accessible process of deliberation. In this case, evidence consists of estimates of state-action values internally generated by the computational process implementing forward planning in the brain, similar to how sensory organs gather evidence by sampling the environment according to Bayesian accounts of perception. In sequential decision tasks, forward planning can generate such estimates by evaluating the future consequences of each action in the current state. However, generating these estimates is typically computationally costly, because it requires evaluating future outcomes in complex and dynamic environments (Sutton et al. 1998; Gershman et al. 2015; Callaway et al. 2022). In analogy to the Bayesian view on perception, agents may therefore combine their evidence, i.e. their internal estimates about future consequences, with prior information about which actions are usually appropriate in a given task. In this view, action selection reflects a balance between a costly state-specific estimate and a stored prior tendency to act in a particular way. This idea has been applied previously in a Bayesian account of the balance between habitual and goal-directed control, in which habitual tendencies were modelled as priors over policies that are balanced against goal-directed information about current outcome contingencies (Schwöbel et al. 2021, 2024). This approach further echoes a recent proposal casting the problem of optimizing planning depth in terms of Bayesian inference (Kuperwajs and Ma 2021). More broadly, it aligns with resource-rational accounts of planning, which postulate that the brain adapts its planning computations to the costs and benefits of using its relatively limited cognitive resources (Gershman et al. 2015; Lai and Gershman 2021; Callaway et al. 2022; Tishby and Polani 2010; Parush et al. 2011).

However, it remains unclear how such prior-like action tendencies are structured in complex sequential decision-making tasks and whether their uncertainty determines reliance on forward planning. In particular, it is unknown whether agents can exploit multiple features of the task to form structured preferences that regulate the influence of planning-derived values. This leads to the central hypothesis of the present study: choice behavior in complex sequential decision-making integrates two sources of action guidance. First, participants may rely on task-specific preferences that summarize regularities learned from task structure or experience. Second, they may use forward planning, as for example typically modelled by reinforcement learning, to compute state-specific action values on the current trial. Assuming a Bayesian approach, we hypothesize that these two sources are integrated by weighting the contribution of forward planning against preferences by their respective uncertainties. This hypothesis implies that on trials where uncertainty about preferences is low, choices should be guided mainly by the preferences, producing faster responses because less forward planning is needed. On trials with less certain preferences, decision making should be more weighted towards forward planning and consequently, response times should be increased due to the increase in computations.

In this paper, we test this hypothesis in a complex sequential decision-making task, using choice model comparisons and response-time analyses. Specifically, we use optimal forward planning as a reference model and ask whether participants’ deviations from this reference are systematic and better explained by including task-specific preferences. We compare this account against several alternative models of how participants may combine planning-derived decision values with potential preferences. Participants’ choices were best captured by a model combining forward planning with task-specific preferences from multiple task features. Response-time analyses converged with this pattern, showing that forward planning conflict slowed responses most when preferences were weak. Together, these findings suggest that costly forward planning is integrated with structured, task-informed preferences that enable faster and less computationally demanding choices when they provide a clear action tendency.

## 2 Results

We tested our hypothesis on a previously published dataset in which participants (N=40) performed a sequential decision-making task with a large state space (Ott et al. 2022b, see Figure 1 and Section 4.1 for the full description of the design). On each trial, participants chose whether to accept or reject an offer worth 1-4 points. Accepting an offer yielded its reward but incurred an energy cost of 1 or 2 points, whereas rejecting yielded no reward but replenished 1 energy point. Costs remained fixed within blocks of four trials (segments), and participants were informed of both the current and the upcoming segment’s cost. Each trial was therefore defined by a combination of four task factors (offer value, energy level, current cost, and future cost), all of which must be integrated to maximize return. This makes the dataset well suited to test our prediction that participants combine forward planning with preferences reflecting the structure of the task.

**Figure 1:**
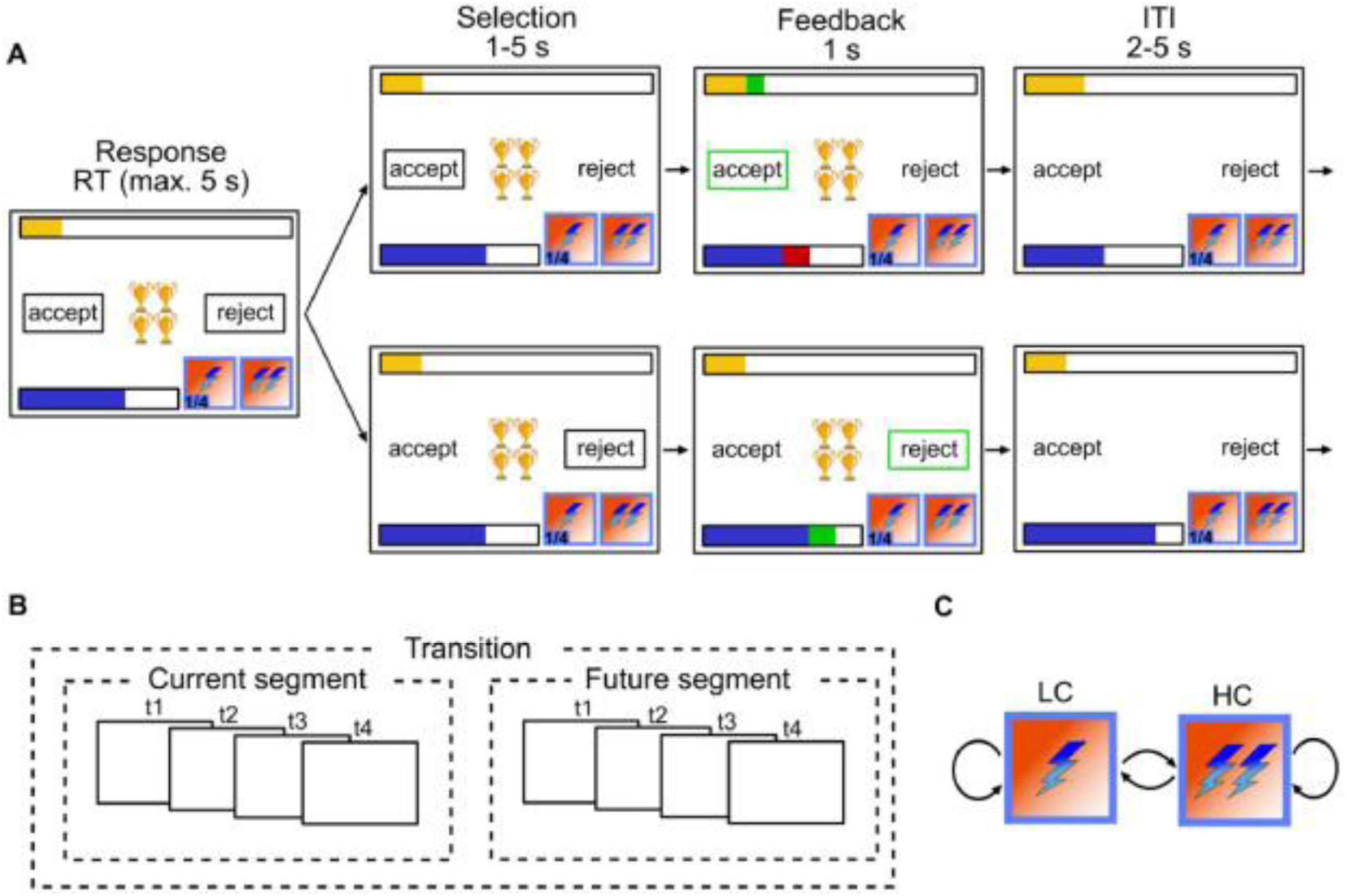
Experimental design: (A) Single trial procedure, each frame represents a step within the trial with duration of each step above. Top row depicts participants accepting; bottom row depicts participants rejecting. In each trial, participants are presented with N golden cups; the number of cups symbolizes the amount of reward. The blue bar represents energy level; yellow bar represents accumulated reward so far, lightning symbols at the bottom right of each frame represent current cost (left) and future cost (right). (B) Between trial costs dependencies. A segment consists of 4 trials and the cost in each segment is fixed. Participants are aware of the cost in the current and future 4 trials but not beyond. (C) Transition structure. Cost can transition from low cost (LC) to high cost (HC), LC to LC, HC to LC and HC to HC. Reproduced from Ott et al. (2022a)

Performance indicates that participants were able to perform the task well. Participants provided responses with an average reaction time (RT) of 0.90s ± 0.26 (s.d.) and failed to give their response before time out in only 0.27% ± 0.56 (s.d.) out of 240 trials. On average, participants gathered a total of 332.43 ± 6.64 points throughout the experiment.

### 2.1 Choice behavior

To test whether and how participants’ choice behavior deviates from optimal planning, we compared four Bayesian hierarchical logistic regression models of participants’ accept/reject choices. All models shared the same planning component: decision values (DV) obtained by backward induction on the task Markov Decision Process (MDP), reflecting the difference in long term value between accepting and rejecting an offer in each state under optimal planning given the specified finite-horizon MDP and terminal-value construction (see Section 4.2). The Planning model contained only DV as a regressor, testing whether deviations are unsystematic noise. The three other models differed in how they accounted for deviations from this benchmark. The Frequency-prior model assumed deviations reflect a recency bias towards recently chosen actions. The Offer-context model, previously found to account for the observed data the best among 21 competing models by Ott et al. (2022a), assumed that participants plan to different extents in extreme versus intermediate offers. Finally, the Preference model, implementing our main hypothesis, combined the planning component with a state-dependent preference score, constructed by assigning weights to the levels of each task factor (offer, energy, current and future costs) and summing them, as well as a modulation term testing whether the influence of planning depends on the entropy of the preference score (see Section 4.3 for a full description of our modeling approach).

Model comparison revealed that participants’ deviations from optimal forward planning were indeed systematic. Replicating Ott et al. (2022a), the Planning model provided the poorest fit (Δ*ELPD_pref_*_−*planning*_=586.28 ± 55.56 s.e., see Figure 2 A), indicating that deviations from optimal behavior are not purely stochastic. Furthermore, our results indicate that participants’ biases do not reflect a frequency prior (Δ*ELPD_pref_*_−*frequency*_ =581.38 ± 55.66), nor a splitting of the task into separate contexts (Δ*ELPD_pref_*_−*context*_=165.49 ± 52.62). Rather, participants exhibit preferences that reflect the task structure, as the Preference model provided the best fit (see Figure 2 A).

**Figure 2:**
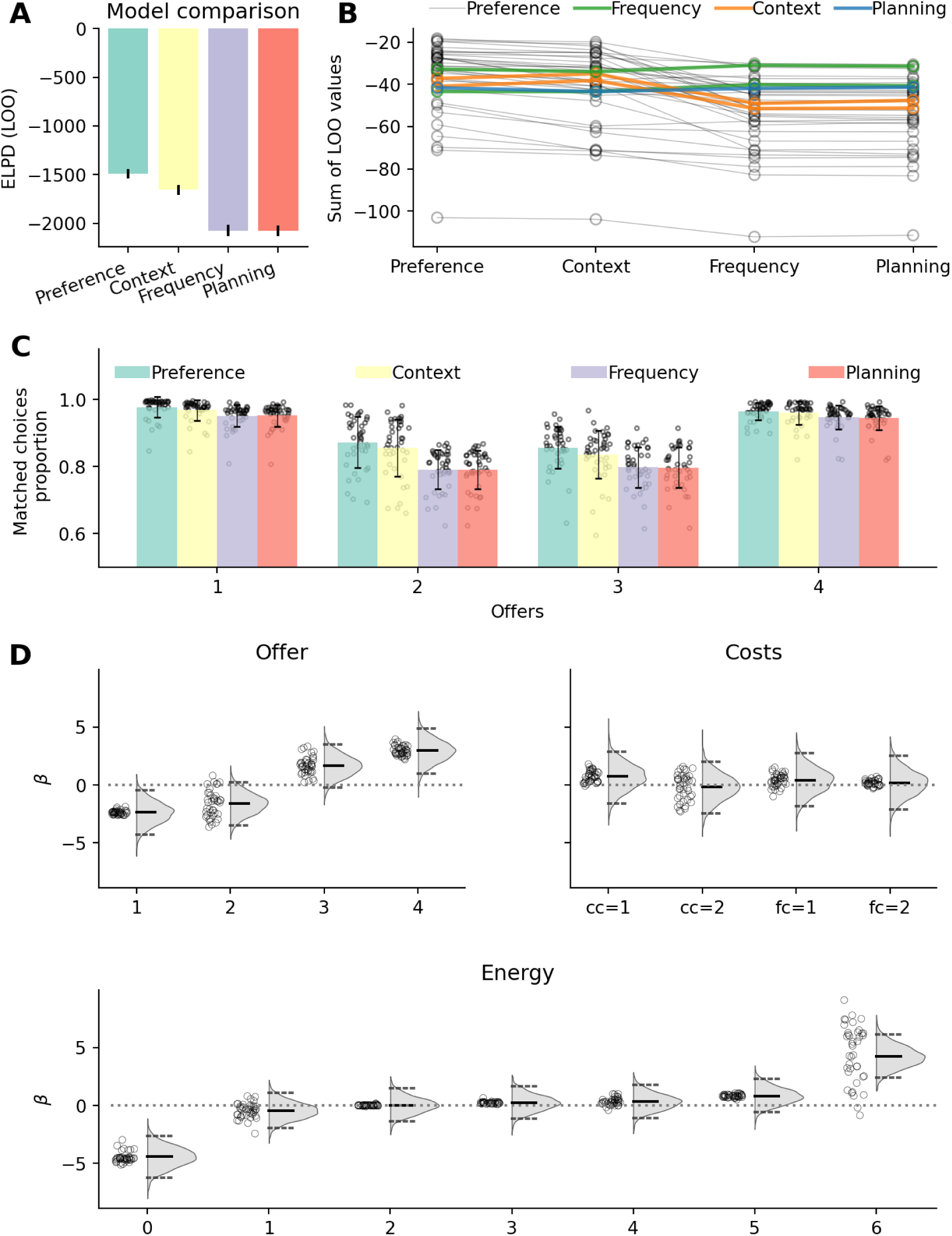
Choice-model comparison and fitted factor-level preference weights (A) Model comparison showing the expected log pointwise predictive density (ELPD) for each model (Preference, Offer-context, Frequency-prior and Planning models). Values closer to 0 indicate better fit. See Table S1 for additional model comparisons results. (B) Participant-wise best-fitting models. Each participant is color-coded according to the model with the highest predictive accuracy. The Preference model provided the best fit for 34 of 40 participants. (C) Proportion of trials in which the model predicted responses match participants’ responses, separately for each model and offer level. Dots represent single participants modelled/observed match proportions. (D) Estimated factor-level preference weights for the Preference model. Panels show weights for offer levels, current and future cost, and energy levels. Distributions show the group-level posterior estimates; central bars indicate medians, and dotted lines indicate the 2.5% and S7.5% quantiles. Dots indicate participant-level mean estimates.

This result is highly consistent across participants: the Preference model provided the best fit for 34 out of 40 participants (see Figure 2 B). The improved fit of the Preference model is reflected in Figure 2 C, showing the proportion of correct choice predictions separately for each model and offer values. Here, the data show that the proportion of matches is on average higher for the preference compared to all other models.

The estimated factor-level preference weights associated with each task factor reveal structured biases in participants’ behavior (see Figure 2 D and Table S2 for the posterior of each fitted parameter). Participants tend to reject low offers (1 and 2) more frequently than optimal forward planning would suggest, while they show a bias towards accepting high offers (3 and 4). These biases are stronger for extreme offers (1 and 4) compared to intermediate ones (2 and 3). The energy state also plays a critical role in shaping participants’ decisions. When energy is at maximum (e = 6), participants almost always accept offers, even when rejecting would be more reward-maximizing. This behavior is intuitive: rejecting offers at max energy does not yield additional energy, so participants prioritize immediate rewards, even at the expense of future rewards. In contrast, when energy is at 0, participants rarely accept offers, even though the planner predicts a small probability of acceptance (∼0.25). This discrepancy arises because the planner calculates DV based on long-term returns, where the difference between accepting and rejecting at energy = 0 is negative but relatively small. However, participants adopt a stricter rejection policy, likely reflecting an aversion to accepting offers that yield no immediate reward. Importantly, energy = 0 occurs in only 17.85% ± 4.02 of trials, so this counterintuitive effect at energy 0 cannot alone explain the success of the Preference model. Finally, costs (both current and future) exert a weaker influence on participants’ behavior, with a mild bias towards accepting offers when costs are low.

We then proceeded to test the central prediction that the influence of forward planning should depend on preference uncertainty, quantified as the entropy of the action tendency implied by the preference score. The Preference model revealed a positive modulation of forward planning by preference entropy (*β_modulation_*=1.26, *CI*_95%_=[0.31, 2.29]), indicating that planning-derived DV had a stronger influence on choices when preferences were more uncertain. Conversely, when preference had lower entropy, choices were less influenced by planning-derived DV and were more strongly guided by the preference score.

This modulation is illustrated in Figure 3 A, where we plotted the fitted probability of accepting as a function of planning-derived DV for different preference scores. Reflecting the main effect of preference score, the logistic curve shifted leftward as the preference score increased. Thus, when preference score favored accepting participants required lower DV to accept an offer. Conversely, when preference score favored rejecting, higher planning-derived DV were required to overcome this preference and lead to acceptance. Reflecting the modulation of preference entropy on DV, the slope of the curve decreases as the preference score deviates further from zero (in either direction) reflecting reduced reliance on forward planning as preferences’ uncertainty decreases. Figure 3 B and C show the observed probability of accepting an offer as a function of DV, separately when preference scores were positive (Figure 3 B) and negative (Figure 3 C). This further shows the effect of preferences on participants’ response choice: participants accept an offer at lower DV when preference scores are positive than when preference scores are negative (see Figure S1 for unbinned participants’ responses and fitted values as a function of DV and preference score).

**Figure 3:**
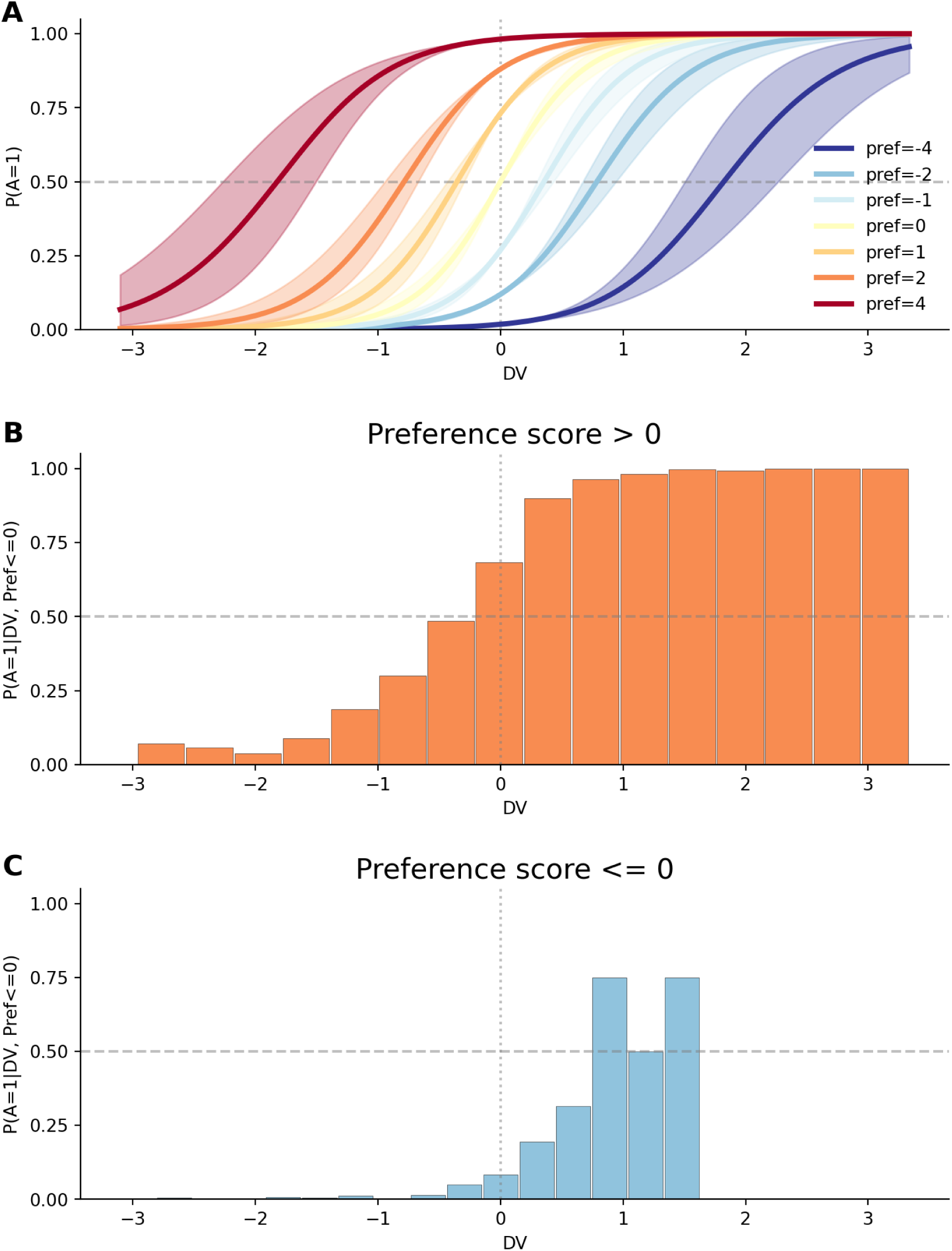
Relationship between preference score and DV. (A) Fitted probability of accepting an offer as a function of DV, separately for preference score of [-4, -2, -1, 0, 1, 2, 4], to highlight the modulation of preference scores on the influence of DV. Lines represent the mean of the posterior distribution; the shaded area is the 95% C.I. (B) Offer acceptance probability (y-axis) binned per DV (x-axis), for trials in which preference score is positive. (C) Same as (B), but for trials in which preference score is negative. Only bins containing observed trials are shown; the different number of bars in panels B and C reflects the unequal distribution of positive- and negative-preference trials across the planning-DV range. This asymmetry arises because negative preference scores are more often associated with states in which forward planning also favors rejection.

We next asked whether the fitted forward planning and preference-related quantities were relevant not only for predicting trial-by-trial choices, but also for explaining individual differences in overall task performance. Specifically, we tested whether overall task performance was related to participants’ reliance on forward planning, their preference entropy, and the extent to which preference uncertainty modulated forward planning. As larger *β_plan_* values indicate stronger reliance on forward planning, we expected participants with larger *β_plan_* to obtain higher returns. In contrast to forward planning, preferences might not necessarily maximize reward, so participants with lower preferences entropy should obtain lower return on average. Finally, larger *β_modulation_* indicate that participants plan more when their preferences are weak (i.e. preferences entropy is larger) and should accordingly obtain larger returns compared to participants who do not engage more strongly in forward planning when their preferences are weak. Accordingly, we predicted that *β_plan_*, preference entropy and *β_modulation_* should be positively correlated with return.

In line with these predictions, we observed a positive correlation between *β_plan_* and return (r=0.58, p<0.001, Figure 4 A), showing that participants with greater fitted sensitivity to planning-derived values tended to obtain higher returns. Furthermore, *β_modulation_* is strongly and positively correlated with return (r=0.66, p<0.001, Figure 4 C). However, contrary to our prediction, we observed a negative though only marginally significant correlation between preference entropy and return (r=-0.29, p=0.072, Figure 4 B). Visual inspection of Figure 4 B revealed that the correlation appears to be driven from the participant with very low return (2.63 s.d. from population mean). When removing this participant, the correlation between preference entropy and return is not marginally significant (r=-0.09, p=0.596), while it remains strongly significant for forward planning (r=0.47, p=0.003) and modulation (r=0.60, p<0.001) when the same subject is removed.

**Figure 4:**
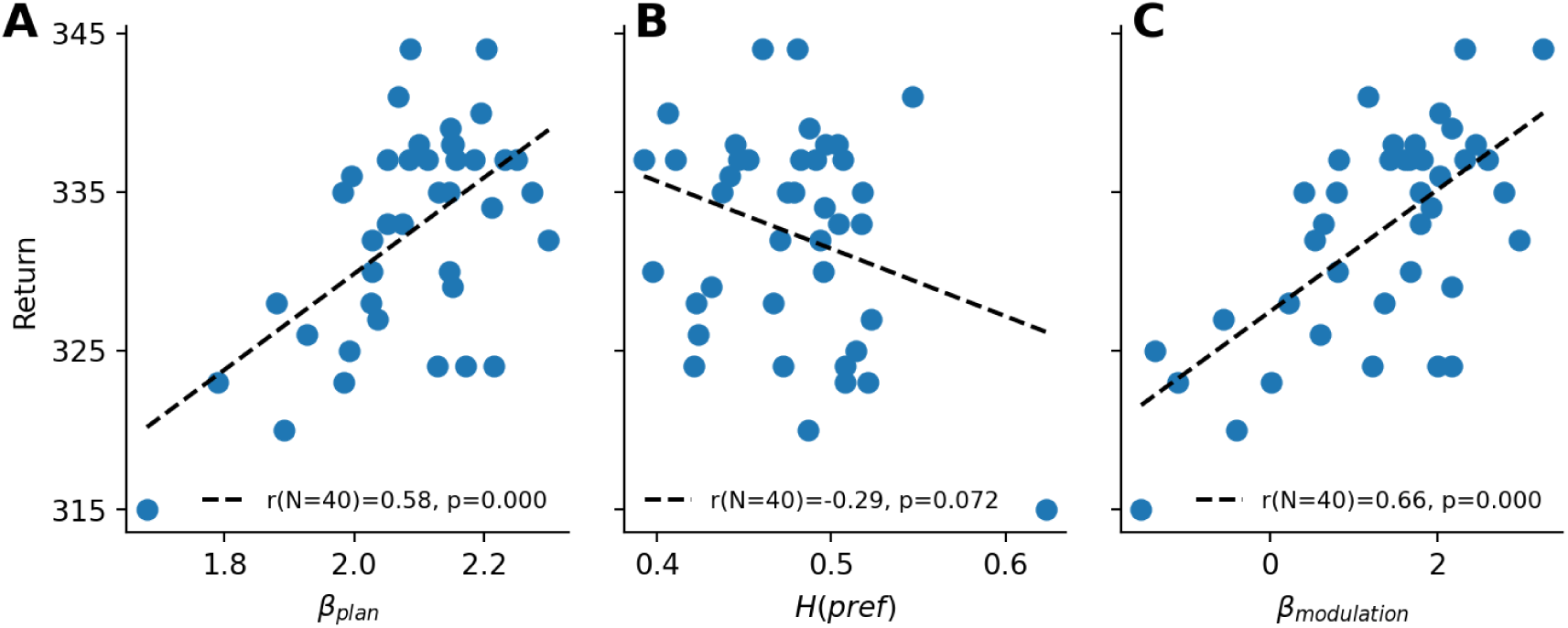
Relationship between fitted parameters and task performance (A) Correlation between participants’ total return and their fitted forward planning sensitivity β_plan_ parameter. Each dot represents one participant. The line shows an ordinary least-squares regression fit, where the legend reports Pearson’s correlation and the corresponding p-value. (B) Correlation between participants’ total return and their average preference entropy. Plotting conventions are the same as in (A). (C) Correlation between participants’ total return and the fitted preference-modulation parameter β_modulation_. Plotting conventions are the same as in (A).

### 2.2 Experimental factors relevance

Our results indicate that participants’ choices are shaped by two sources of information: planning-derived values and task-specific preference scores. These preference scores are not a single global bias. Instead, under the Preference model, state-dependent preferences reflect a trial-wise weighted sum of preferences associated with different factors of the task such as the offer, the current cost context, and upcoming transitions. Importantly, not all features of the task are weighted equally. Figure 2 D shows that the parameters associated with offers and energy weigh strongly in participants’ preference scores compared to costs (both current and future). The fitted factor-level preference weights already suggested that participants’ preference scores were structured primarily by offer value and energy. We further confirmed this pattern from a complementary model-comparison perspective. To do so, we compared the full Preference model with reduced models in which the factor-level preference was removed for one task factor at a time (see Section 4.5 for a description). This effectively tests how decision-making changes when participants are assumed to ignore one specific source of task information, such as offer value, action costs, or future transitions.

Figure 5 A shows the change in model fit when each preference factor was removed separately from the full Preference model. Each reduced model omitted the preference term for one experimental factor while retaining the remaining preference terms. Larger values indicate that the specific factor is more relevant for the full model’s fit. We found that offer (Δ*ELPD_full_*_−*offer*_=137.36 ± 54.44 s.e.), followed by energy (Δ*ELPD_full_*_−*energy*_ =133.97 ± 51.57 s.e.), has a much larger impact on participants’ final responses, whereas costs-related preferences have a smaller impact on the overall fit (Δ*ELPD_full_*_−*CC*_ =47.02 ± 49.89 s.e. and Δ*ELPD_full_*_−*FC*_=27.35 ± 48.62 s.e.).

**Figure 5:**
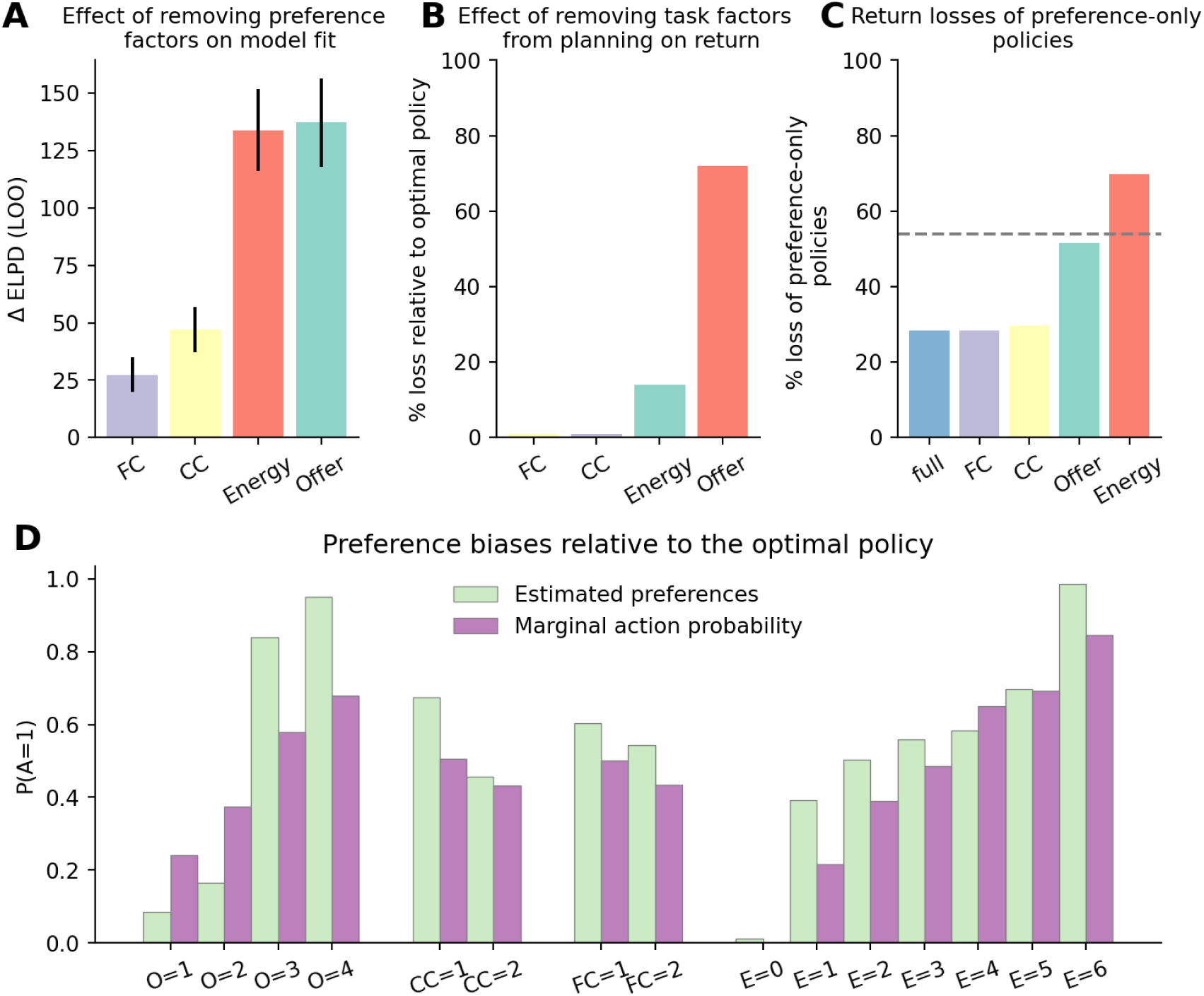
Contribution of task factors to preference-based choice and performance. (A) Predictive contribution of individual preference factors. Difference in fit between the full Preference model and models in which each of the experimental factors was removed from preference score estimations in turn. Bars show ΔELPD, computed as the ELPD of the full model minus the ELPD of the corresponding reduced model. Values closer to 0 indicate that a factor does not lead to a strong improvement of the model fit (i.e. the factor weighs less in participants’ final decision). (B) Loss in return by ignoring individual task factors during forward planning (0 indicates no loss compared to the return under the optimal policy derived from the full MDP). Bars show the reward sensitivity of each task factor: larger values indicate that the omitted factor was more important for reward-maximizing planning. (C) Return loss of policies derived from the preference score of the full and reduced Preference models, in which the preference terms for one task factor were omitted in turn. For each model (full and reduced), we derived a policy from the preference score alone and calculated the expected return under such a policy, relative to the optimal policy. The y-axis represents the loss in relative return; 0 indicates that a preference policy performs exactly as well as the optimal policy. The horizonal dashed line represents the return associated with a fully stochastic policy (i.e. equal probability of each action in each state). (D) Per factor-levels accept tendencies in the fitted preferences and in the optimal policy. For each level of each task factor, we calculated the accept tendency implied by the fitted factor-level preference weight and the average accept tendency of the optimal policy in states containing that feature level, after averaging over all other task factors. Similar profiles indicate that participants’ preferences capture task regularities present in the optimal policy; deviations indicate feature-specific biases.

We further sought to determine whether the fitted preferences could serve as adaptive biases when forward planning is absent. As a first step, we quantified how strongly optimal return depends on each task factor. To do so, we derived policies from reduced forward planning models in which one task factor was ignored and computed their return loss relative to the optimal policy (Figure 5 B, see Section 4.5 for more details). Our results show that ignoring current and future costs when planning leads only to minimal losses (0.78% and 0.98% respectively), while ignoring the offer level leads to a 13.94% loss. The biggest loss is incurred when ignoring energy levels. This leads to a loss in returns of 72.04%, which is even larger than a purely stochastic policy (i.e., equal probability of accepting and rejecting an offer on each trial). This indicates that reward maximization under forward planning is highly sensitive to energy. This large loss reflects the structure of the task. A policy that ignores energy cannot adapt choices to the current energy budget. As a result, it can deplete energy in low-energy states as its behavior is driven by offer levels. The large reward sensitivity of energy is not mirrored in Figure 5 A because that analysis measures the additional predictive contribution of preference terms after accounting for planning-derived decision-values. Energy is already strongly reflected by the forward planning component, so removing energy-related preference terms has a smaller effect on model fit than its relevance for optimal return would suggest.

Further, we asked whether the fitted preferences could support adaptive choice when forward planning is absent. If preferences capture useful task structure, policies derived from the preference score alone should be suboptimal but still produce less return loss than a stochastic policy. In addition, removing task factors that strongly shape participants’ preferences should reduce the return of these preference policies. To quantify this, we computed the expected return loss of policies derived from the preference score alone (see Section 4.5). The full preference policy produced a return loss of 28.34%, indicating that preferences did not fully replace forward planning, see Figure 5 C. However, this loss was smaller than the loss of a stochastic policy (dashed line), showing that the fitted preferences captured task structure that was relevant for return maximization (Figure 5 C). Removing current or future cost from the preference policy produced little additional loss relative to the full preference policy (29.72 and 28.38 respectively). In contrast, removing offer value led to a substantial loss in return of 51.52%. The largest loss occurred when energy was removed (69.80% loss), with performance falling below that of the stochastic policy. This suggests that an energy-blind preference policy is not merely less informative but can become systematically maladaptive by accepting attractive offers without regard to the current energy budget, making the agent more likely to run out of energy than under a stochastic policy.

Together, these results suggest that participants’ preferences were not arbitrary biases, but adaptive approximations to optimal choice. They supported relatively good performance even in the absence of forward planning (loss of 28.34% from optimal returns) and provide a useful first-pass policy that can support good performance when forward planning is limited but can be further refined by planning-derived DV when more computation is available.

How do participants derive such adaptive preferences? A potential explanation is that participants have access to some approximation of the marginal action probability associated with each factor of the task under the optimal policy (for example, how often should an offer of 1 be accepted, across all energy and costs levels). To explore whether this might be contributing to an underlying mechanism for learning participants’ preferences, we computed the factor-dependent marginal action probability under the optimal policy. Figure 5 D plots the factor-level preferences parameters side-by-side with the marginal action probability, showing that participants’ preferences reflect relatively closely the factor-level specific marginal action probability of the optimal policy, with some notable deviations. For offer levels 1 and 2, preferences are lower than marginal action probability, and higher when for offer levels 3 and 4. Across all other tasks levels, participants’ preferences are larger than marginal action probability (except for energy=4), suggestive of an optimistic bias (i.e. a tendency to accept offers more often than participants should). This optimistic bias might be afforded by the gain in energy associated with the tendency to reject low offers. Nonetheless, the overall similarities between both variables suggest that these marginal action tendencies may have contributed to the formation of participants’ task structured preferences.

### 2.3 Reaction times

Under the proposed hypothesis, participants’ reliance on forward planning should depend on preference entropy. Low preferences entropy should reduce the need for forward planning and therefore lead to faster responses, whereas high preferences entropy should increase reliance on forward planning and thereby lead to slower responses. Previous studies have shown that RT is positively correlated with forward planning conflict, indicating that less forward planning is required when the value associated with each action is more clearly distinct (Ott et al. 2022a; Gluth et al. 2012; Lee and Daunizeau 2021). We therefore expected preference scores to attenuate the effect of forward planning conflict on RT: when participants’ preferences provide a clear action tendency, forward planning conflict should have a weaker influence on response times. To test this prediction, we modelled participants’ RT using planning conflict (−|DV|), i.e. the negative absolute decision value, as well as the preference score entropy estimated from the choice behavior model, and the interaction between the two (see Section 4.6 for a description of the model).

Indeed, we found that participants’ RT was influenced by (i) forward planning conflict (*β_conflict_* =0.11, *CI*_95%_=[0.08, 0.14]), (ii) preference entropy (*β_pref_* =0.45, *CI*_95%_=[0.35, 0.54]) and (iii) the modulation effect between the two (*β_modulation_*=0.11, *CI*_95%_=[0.06, 0.15], see Figure 6 A). This result extends previous findings showing that participants do not rely on forward planning to the same extent across all states of the task. Our results suggest that rather than splitting the task in independent contexts and engaging in forward planning to a different extent across those (Ott et al. 2022a), the differential amount of forward planning across states instead reflects the fact that participants engage in forward planning more when preferences are weak. Specifically, as can be seen in Figure 6 B, forward planning depends less on forward planning conflict when preferences entropy is low (blue line), but more so when preferences entropy is larger (orange and green lines).

**Figure 6:**
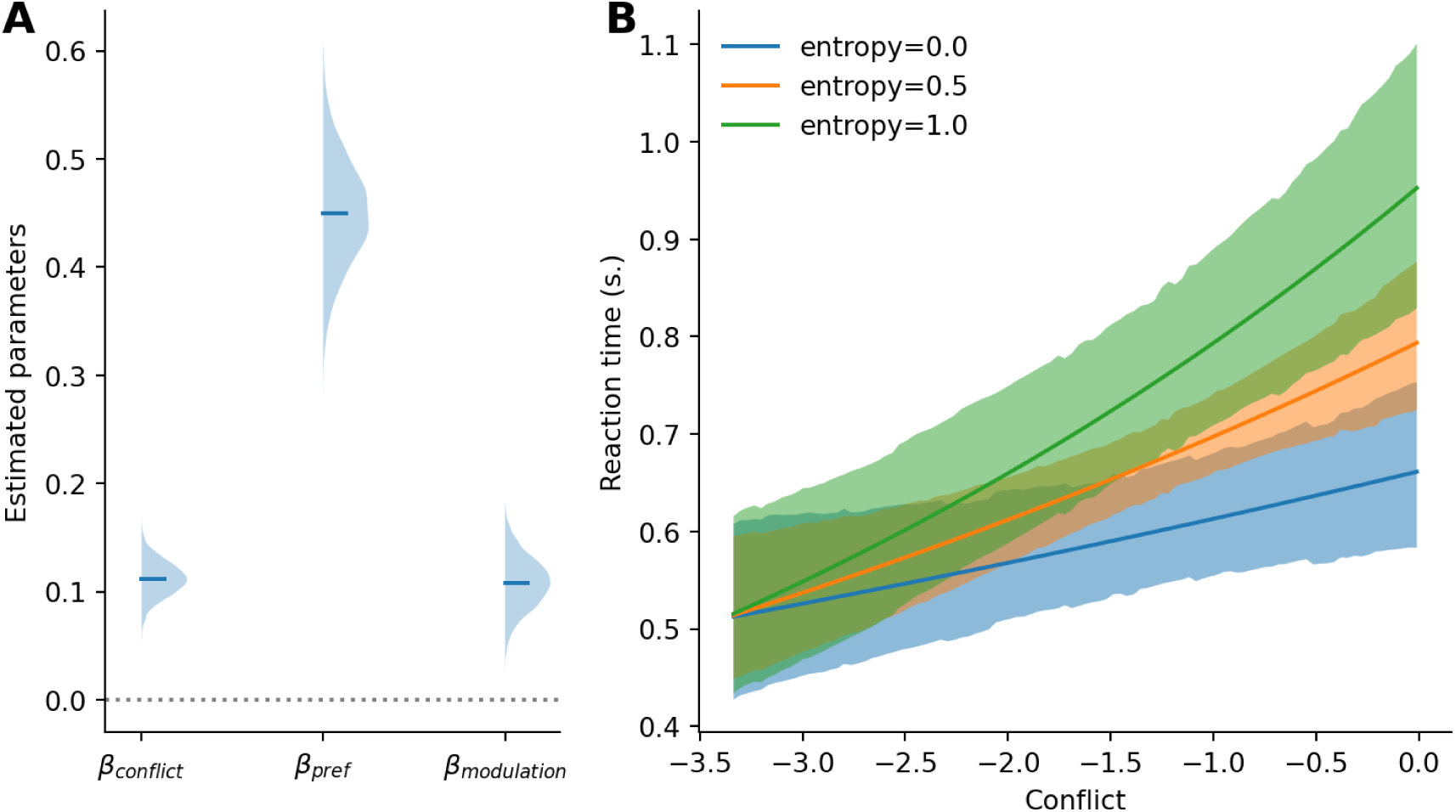
RT evidence for preference-modulated forward planning. (A) Posterior distributions of the fitted response-time effects, including forward planning conflict, preference entropy, and their interaction. (B) Posterior predictive curves depicting the predicted RT as a function of forward planning conflict, separately for preference entropy of 0, 0.5 and 1.0.

Together, these response-time results converge with the choice-model findings. Participants were slowest when planning-derived DV were in conflict, and task-specific preferences had high entropy, indicating stronger reliance on forward planning under preference uncertainty (green line). Conversely, when preferences provided a clear action tendency, forward planning conflict had a weaker effect on response times (blue line), suggesting reduced reliance on planning-derived values.

## 3 Discussion

We found that participants’ deviations from an optimal forward planning reference were systematically organized by task-specific preferences, and that these preferences regulated the influence of planning-derived decision values on choice. This pattern was not well explained by unsystematic errors, recent action-history effects, or the offer-context biases identified in previous work. Instead, participants’ choices were best explained by a model that combined planning-derived decision values with task-specific preferences over offers, energy, and costs. Crucially, these preferences did not act only as additive choice biases. Planning-derived values had the strongest influence on choice when preference uncertainty was high, whereas lower preference uncertainty was associated with reduced planning-related computation, as evidenced by participants’ RT. Additional policy analyses indicated that these preferences preserved reward-relevant task structure and supported substantially higher returns than stochastic choice even without forward planning, although they remained below the optimal forward planning policy. Feature-wise analyses showed that these preferences captured reward-relevant structures of the task. Together, these results suggest that participants used task-specific preferences as efficient approximations to reward-maximizing choice, while relying more strongly on forward planning when these preferences did not provide a clear action tendency.

This finding reframes task-factor effects in sequential decision-making. Rather than treating offer, energy, and cost effects as separate descriptive biases, the present account interprets them as structured action tendencies that interact with planning-derived values. Computational models of sequential decision-making often include additional parameters capturing response biases, perseveration, or other deviations from value-based choice (Ott et al. 2022a, 2020; Ho et al. 2022; Huys et al. 2012; Schwartenbeck et al. 2015). However, these terms are typically low-dimensional and are not usually decomposed according to the structure of the task. Conversely, factorial analyses of experimental data can quantify how different task factors influence behavior or neural responses (Barr et al. 2013; Baayen and Milin 2010; Ashburner et al. 2014; Singmann and Kellen 2019), but they do not by themselves specify how these effects relate to forward planning. By combining these approaches, our model treats task-factor effects as structured action tendencies that can guide choices and modulate reliance on planning-derived values.

Our approach was motivated by Bayesian accounts of perception and sensorimotor control, in which current evidence is integrated with prior information according to their relative uncertainty (Knill and Pouget 2004; Lee and Mumford 2003; Rao and Ballard 1999; Clark 2013; Friston 2010; Körding and Wolpert 2004). An intuitive interpretation of our results is that task-specific preferences play the role of priors, whereas planning-derived values provide the likelihood-like, state-specific evidence about which action will lead to better future outcomes, and the entropy of the priors determines the extent of forward planning. In our task, this interpretation is particularly evident in how preferences guide behavior in different states. In states where the relevant features jointly imply a clear action tendency (e.g. high offer, low cost and high energy), preferences alone may be sufficient to support efficient decision-making, requiring minimal forward planning. In other states, different features support opposing actions. For instance, a moderate offer may favor rejection, whereas low cost and high energy may favor acceptance. Such opposing factor-level tendencies produce a preference score closer to zero and therefore higher preference uncertainty. In such cases, forward planning can help resolve the remaining ambiguity by evaluating the future consequences of the available actions.

Our model resembles the policy compression framework, under which forward planning values are integrated with action priors in a Bayesian fashion (Lai and Gershman 2021). Importantly, this framework assumes state-independent priors, reflecting the marginal action probability under the current policy. While such priors work well in non-sequential decision-making tasks or in tasks with limited downstream consequences (Gershman, 2020; Lai and Gershman 2024; Bari and Gershman 2023; Liu and Gershman 2025; Liu, S. et al. 2025; Lai et al. 2022; Gershman and Lak 2025), they may be insufficient in more complex, sequential decision-making tasks, where action–state combinations have complex downstream implications. Our results suggest that participants rely on state-dependent preferences in such tasks. The factorized preference components strike a balance between fully state-specific priors, which would be overly costly in terms of memory and completely state-independent action priors that would be too coarse-grained to be useful in complex tasks.

Together, these findings provide a potential refinement of the context-dependent planning effects reported by Ott et al. (2022a). In that study, participants showed a greater propensity to engage in forward planning for intermediate offers (2 and 3) than for extreme offers (1 and 4), with model comparisons supporting a coarse partition of the state space into these offer contexts. Our results suggest a more general principle: intermediate offers are associated with less decisive preference-based action tendencies, and therefore with greater reliance on planning-derived values. Thus, rather than varying only across predefined offer contexts, planning-related control may vary with preference uncertainty over the relevant task factors, including offer, energy, current cost, and future cost.

Our results contribute to the debate about whether deviations from optimality reflect systematic errors or adaptive heuristics. Across psychology and perceptual decision-making, deviations from normative models have often been described as systematic errors (Tversky and Kahneman 1974; Gilovich and Griffin 2002; Berthet and De Gardelle 2023). In contrast, biases can also be interpreted as adaptive heuristics that provide computationally inexpensive shortcuts and support effective, though not necessarily optimal, solutions in complex tasks (Gigerenzer et al. 2000; Tsetsos et al. 2016; Gigerenzer and Brighton 2009). In line with an adaptive-heuristic interpretation, our findings suggest that preferences were not arbitrary residuals around an optimal-planning process. Rather, they appeared to capture structured action tendencies derived from task features, which supported above-stochastic performance even in the absence of forward planning. Deviations from optimal forward planning may partly reflect inexpensive, task-specific action tendencies that provide a useful first-pass guide, while being complemented by forward planning when further evaluation is needed. This pattern suggests that preference-based guidance and planning-derived evaluation interact within a single decision process, rather than reflecting a simple opposition between heuristic bias and optimal forward planning.

Importantly, while the Preference model suggests that task-specific preferences can function like prior action tendencies, the model is rather descriptive in the sense that it does not establish that the fitted preferences are priors in a formal Bayesian sense. In addition, the model implies that the preferences were stable during the modelled task phase and does not capture updating of these preferences. The similarity between preferences and factor levels-dependent marginal action probabilities (see Figure 5 D) suggests that preferences might arise by keeping track of actions performed at each level of each factor under an approximately optimal policy. For example, this may have happened in the initial training session (Ruge and Wolfensteller 2010). Furthermore, general schemas from encounters with similar naturally occurring problems in the environment (budgeting decisions in everyday life), or explicit instructions highlighting relevant features might also play a role in fast learning of preferences. A key question for future research is whether such preferences function as state-dependent action priors, and how agents determine the task features that are relevant for constructing and learning these priors, especially in real-world settings where task structure is not always explicit.

While our model uses backward induction to derive state-action values, this does not imply that the brain implements forward planning in this way. Biologically plausible forward planning may rely on approximate algorithms such as sampling, partial tree search and approximate value computations (Sutton et al. 1998; Silver et al. 2016; Huys et al. 2012; Schwöbel et al. 2021). We are not committed to any one of these alternatives, as our main interest in the paper was to investigate the structure underlying participants’ deviations from forward planning. The observed interaction between RT, preferences entropy and forward planning conflict suggests that whichever algorithm implements action value estimation in the brain, the extent of planning-related computation can be adaptively adjusted to obtain estimates of different precision, depending on preferences entropy. Developing RL or active inference agents equipped with such structured priors and testing their predictions against human data may be important for understanding the emergence and utility of efficient, adaptive decision strategies in complex environments.

The present account makes several testable predictions about when preference-based guidance should dominate. Reliance on preferences should increase when task features provide a clear action tendency, when forward planning time is limited, or when the marginal value of additional computation is low. Conversely, reliance on forward planning should increase when task features provide conflicting action tendencies, when future consequences are difficult to infer from local features alone, or when the cost of an error is high. Future experiments could test these predictions by manipulating factors such as time pressure, training, instructions, and task familiarity, planning horizon, task volatility, or the reliability of feature-based cues. Such manipulations could help disentangle the sources of task-specific preferences and clarify how structured priors are formed, adapted and integrated with forward planning.

## 4 Methods

### 4.1 Data

The data used in this study were published by Ott and colleagues (2022a). In this study, 40 participants (22 women, mean age=24.4, SD=4.6) took part in a sequential decision-making task (Ott et al. 2022a, 2022b). The study was approved by the Institutional Review Board of the Technische Universität Dresden and conducted in accordance with ethical standards of the Declaration of Helsinki.

Figure 1 shows a schematic representation of the task. Participants were instructed to gather as many points as possible throughout the task. In each trial, they were presented with an offer associated with a reward of either 1, 2, 3 or 4 points, which they could either accept or reject. To accept the offer in a trial, participants had to pay an energy cost of either 1 or 2 energy points (low cost, LC and high cost, HC respectively). When they rejected the offer, they gained 1 energy point. Participants’ energy level was capped at 6, so their energy could range from 0 (depleted energy) to 6 (max energy). Offers varied in each trial following a uniform distribution. Costs remained fixed for 4 consecutive trials, which we refer to as a segment (see Figure 1 B), and participants were informed of the price of the current segment and the next segment. The cost of the next segment was pseudorandomized to equate the probability of each transition (from current HC or LC to future HC or LC, see Figure 1 C) and ensure that each transition occurred 15 times. When participants accepted an offer that they could not afford (current energy level below current cost), no reward was awarded and a warning was displayed.

Accordingly, each trial was defined by several task factors: the current offer, the participant’s current energy level, the current cost of accepting, and the cost in the upcoming segment. Participants had to decide whether to accept or reject the offer under the specific combination of these factors. Because accepting uses up energy while rejecting replenished it, each choice also changed the state in which the participant entered subsequent trials. Maximizing overall return was therefore non-trivial and required considering the future consequences of current choices.

At the start of each trial, a fixation cross was presented for 0.5 s in the middle of the screen. Then, an offer appeared represented by the number of “cups” presented in the middle of the screen, and choice options were presented on each side of the screen, surrounded by a black frame. Participants had 5 s to provide an answer, by selecting either option (accept or reject), in which case the frame around the chosen option disappeared. Following their choice, the resulting change in energy and points was displayed (green indicating an increase in point and/or energy, red indicating a decrease in energy) for 1s. The trial ended with an inter-trial interval (ITI) ranging from 2 to 5 s (uniformly sampled, rounded to the first decimal) in which the offer disappeared, and unframed choice options remained on screen. If participants took longer than 5 s to provide their answer, a warning message was displayed and the next trial started.

Following their choice, the resulting change in energy and points was displayed (green indicating an increase in point and/or energy, red indicating a decrease in energy) for 1s. The trial ended with an inter-trial interval (ITI) ranging from 2 to 5 s (uniformly sampled, rounded to the first decimal) in which the offer disappeared, and unframed choice options remained on screen. If participants took longer than 5 s to provide their answer, a warning message was displayed and the next trial started.

To familiarize themselves with the task, participants conducted a training session of 144 trials, corresponding to 36 segments. Participants then completed 240 trials (60 segments) which are included in subsequent analyses. Participants started with an energy budget of 3, and their reward points were accumulated throughout the entire experiment. The trials sequence was pseudorandomized, such that each costs transitions (LC to LC, LC to HC, HC to LC and HC to HC, see Figure 1 C) occurred equally often within the training and test sessions, resulting in 9 repetitions of each transition in the training session and 15 in the experiment. In addition, the offer sequence was selected such that the frequency of each offer was matched across costs transitions. The sequence of costs’ transitions and offers was kept the same across participants. Participants were offered two to three minutes breaks every 20 segments. The test sessions lasted around 40 min.

### 4.2 Forward planning component

To model optimal behavior in the task, we operationalized our task as a Markov Decision Process (MDP) with the following tuples:

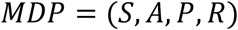

Where *S* represents all possible states in our task, where each state *s* consists of a combination of the experimental variables in our task: *S* = *E* × *O* × *CC* × *FC* × *T*, with energy *E* = 0,1, … ,6, offer *O* = 1,2,3,4, current segment cost *CC* = 1,2, future segment cost *FC* = 1,2 and trial *T* = 1,2, … ,13. Time steps (i.e. trials) had to be incorporated in the state space to account for the cost being trial dependent. *A* is the set of possible actions in our task (*accept* = 1, *reject* = 0), *P* is the transitional probability from a given state to all other states (*P* = *P*_∀*s*∈*S*_ (*s*^′^|*s*, *a*)), and *R* is the reward function characterizing the reward participants can obtain in a given state for each action (*R* = *R*_∀*s*∈*S*_(*s*|*a*)).

While optimal behavior would require considering outcomes up until the end of the task, we modelled forward planning over a limited horizon, following the approach used by Ott et al. (2022a). Participants had explicit information about the current and upcoming segment, corresponding to an eight-trial horizon. Solving the MDP only over these eight trials would implicitly treat the task as ending after the upcoming segment, leading the optimal policy to spend all remaining energy by the end of that horizon. To avoid this terminal artefact, we included one additional segment beyond the known horizon. Thus, action values were computed over a twelve-trial planning horizon, with a terminal value set after the twelfth trial, yielding 13 time points in the dynamic-programming solution. In the first 8 trials, the probability of the next state reflects the offer-related stochasticity, while costs are known. For trials beyond the known horizon, the uncertainty associated with the offer is compounded by the uncertainty related to the next cost transition (see Ott et al. 2022a for more details).

We applied backward induction to compute the long-term value associated with each action in any given state (i.e. the *Q* function Sutton et al. 1998). Participants maximizing return should choose the action with the largest value. As there were only two possible actions in our task (accept or reject), we computed decision values (DV) as:

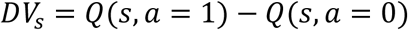

In states where the DV are negative, participants should reject, and in states where DV are positive, participants should accept the offer.

### 4.3 Modelling of participants responses

Participants responses were modelled using a logistic regression:

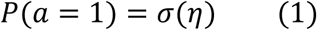

where 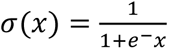 denotes the logistic sigmoid.

Participants’ responses were modelled using Bayesian hierarchical logistic regressions that estimate the probability of accepting the current offer. We compared four models that differed in which sources of information were allowed to influence this probability. All models included planning-derived DV as a benchmark estimate of optimal action value. The models differed in how they accounted for systematic deviations from this forward planning benchmark: as unstructured noise, recent action frequencies, the Offer-context structure identified in the original analysis by Ott et al. (2022a), or task-specific preferences. This comparison builds on previous work by including the best-fitting model among the 21 models tested by Ott et al. (2022a) as a strong reference model. We then asked whether a factorized preference account could explain participants’ choices better than this previous account. Additional parametrizations, reduced models, and robustness checks are described in the model-comparison section below and reported in the supplementary material. Formally, these differences were implemented by changing the predictors included in the linear predictor *η*. We describe the models in turn, starting with the planning model and then adding increasingly structured accounts of participants’ deviations from optimal forward planning.

#### 4.3.1 Planning model

To establish a benchmark, we first asked whether participants’ choices could be explained by optimal forward planning alone. In this pure Planning model, participants’ responses were modelled only as a function of the planning-derived DV:

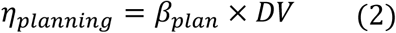

This model captures the possibility that participants’ deviations from optimal forward planning are unsystematic. Under this account, alternative models with additional regressors should not yield better predictive accuracy once model complexity is taken into account. Any improvement of the subsequent models would therefore indicate that deviations from the forward planning benchmark contain systematic structure.

#### 4.3.2 Frequency-prior model

Alternatively, participants’ deviations from optimality might reflect a recency bias, whereby choice on a given trial is biased towards the most frequent action in recent history. Recent accounts of forward planning consider that priors consist of a state independent recency bias (Schwöbel et al. 2021, 2024). Similarly, under the policy compression framework, the optimal policy integrates forward planning and marginal action probability across states (Gershman 2020; Lai and Gershman 2021). To test whether in our task participants’ responses are biased towards most frequent action taken in previous trials, and approximate the policy compression framework, we fitted a Frequency-prior model, which models participants’ responses as a function of DV combined with a recency bias:

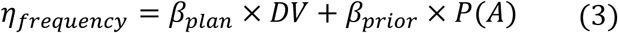

We computed *P*(*A*)*_t_* in each trial *t* using a linearly decaying moving average going back *k* = 20 trials, assuming temporally discounted effects of past actions (see Figure S3 for exploration of the fit at different *k*):

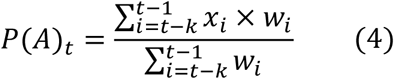

Where *x_i_* denotes the action taken on trial i coded as accept or reject, and *w* = [1,2, … , *k*] so that the weights increase linearly from 1 for the oldest included trial to *k* for the most recent trial.

#### 4.3.3 Offer-context model

In addition, we used the best-fitting model (among 21 models) from Ott et al. (2022a), to test whether participants treat states clusters as separate contexts in which they plan to different extents. We refer to this model as the Offer-context model. Here, the forward planning component (DV) is split between extreme and intermediate offers (offers [1, 4] and [2, 3] respectively) to reflect the fact that participants treat those as separate contexts, with different control requirements. In addition, the model comprises offer specific and energy related flexible biases regressors:

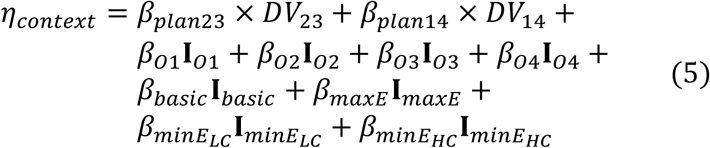

**I** indicate dummy regressors encoding specific experimental conditions: **I***_basic_* encodes trials where energy is sufficient to accept the offer and inferior to 6, **I***_maxE_* encodes trials where energy is equal to 6, **I***_minELC_* for trials where energy is too low to accept the offer when cost is low, **I***_minEHC_* for trials where energy is too low to accept the offer when cost is high, **I***_O_*_1−4_ for trials with offer 1-4 where energy is sufficient to accept. Following Ott et al. (2022a), these regressors meant to account for edge cases within the state space were not included in the Frequency-prior model as it was aimed to test whether participants’ biases from optimal behavior can be uniquely explained by a frequency bias. These were also not included in the Preference model below, as these regressors consist of a linear combination of the preference factors of the model and would therefore be redundant.

#### 4.3.4 Preference model

Finally, we developed a task-specific preference model (Preference model for short) to test whether participants’ choices reflect the integration of forward planning with task-specific preferences. The model contains three components. First, as in all other models, choices depend on the planning-derived DV. Second, choices depend on a state-dependent, task-specific preference score. This preference score is constructed by assigning preference weights to the levels of the main task factors namely offer, energy, current cost, and future cost. We refer to these as factor-level preferences (i.e., the preference weight associated with a given level of a given factor). The score is then computed as the sum of the corresponding factor-level preferences for the current state. Third, the model includes a modulation term that tests whether the influence of forward planning depends on the uncertainty of the preference score. We quantified this uncertainty as the entropy of the preference score (preference entropy). The model therefore tests the prediction that participants rely more on forward planning when their preferences are weak or uncertain:

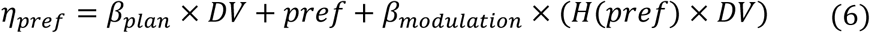

Where:

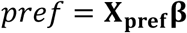

And *H*(*pref*) is the entropy of the preference score:

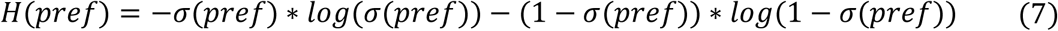

and *σ*(*x*) is the logistic sigmoid defined above.

Participants’ factor-level preferences are estimated as latent variables. **X_pref_** is a [*M* × *N*] (M=measurements, N=experimental levels) matrix dummy coding each factor of the experimental setup (energy, offer, current and future costs) and **β** is a vector of weights estimated for each level of each factor.

### 4.4 Model fitting and comparison

Models were fitted to participants’ response data using Bayesian hierarchical logistic regressions implemented using PYMC (Abril-Pla et al. 2023) and Bambi (Capretto et al. 2022). We excluded trials where participants did not provide a response or if RT exceeded the response window (5 s). Across models, the following parameters were modelled hierarchically, with separate participant-level estimates drawn from group-level distributions: *β_plan_, β_plan23_, β_plan14_, β_pref_, β_modulation_, β_basic_, β_O1_, β_O1_, β_O2_, β_O3_, β_O4_*, while the remaining parameters were fixed across participants. For all parameters, we used weakly informative hyperprior distributions *μ* ∼ N(0,2) and *σ* ∼ *Halfnormal*(0,2). All models were fitted using 4 chains of 2000 samples each (1000 warmups), resulting in 4000 samples in total. We report the mean of the posterior and the 95% credible interval for the estimated parameters in the text.

We used Pareto-smoothed importance sampling to approximate leave-one-out cross-validation (PSIS-LOO, Vehtari et al. 2017) to estimate the expected log pointwise predictive density (ELPD) which we used to compare the fit of these different models accounting for model complexity. Furthermore, we performed model comparison within each single subject to estimate the robustness of the best-fitting model in our population sample, by computing the sum of the pointwise predictive accuracy for each participant and model, yielding a score for each participant and model.

Note that each of the four main models was compared against alternative parameterizations to ensure the robustness of our findings and rule out alternative explanations. For example, the Frequency-prior model was tested with varying temporal integration windows (see Figure S3 and Table S4), and the Offer-context model was selected from a previous study (Ott et al. 2022a) after comparison with 21 alternatives. Similarly, reduced versions of the Preference model were tested against the full model by removing each component (forward planning, preference and modulation) as well as individual task factors (or a total of 7 alternative preference models, see Figure S2 and Table S3). For clarity, we focus on the four best-fitting models (Planning, Frequency prior, Offer-context, and Preference models) in the main text, with comparisons to all alternatives detailed in the supplementary material.

### 4.5 Experimental factors relevance analysis

The Preference model estimates factor-level preference weights for each task factor (energy, offer, current and future costs), revealing that participants’ deviations from optimal forward planning are more strongly driven by some factors of the task (energy and offers) than others (current and future costs). To quantify the behavioral contribution of each factor, we compared the full Preference model to reduced models in which all factor-level preference regressors from one task factor were removed. A large decrease in predictive accuracy after removing a factor indicates that preferences associated with this factor explain a substantial part of participants’ choices beyond the planning-derived values.

The observation that participants’ deviation from optimal forward planning is systematically influenced by specific task factors raises the question of whether preferences constitute unhelpful biases (suboptimal tendencies without adaptive value in the task) or whether they reflect a structured yet simplified solution to the task. In other words, could participants’ preferences constitute a low-dimensional decomposition of an approximation of the optimal policy, providing a cheap alternative to guide behavior in the absence of forward planning? To test this hypothesis, we conducted a series of analyses to test whether weights of the factors in participants’ preferences relate to the structure of the task.

First, we asked whether the factors that are strongly weighted in participants’ preference scores are more relevant for reward maximization. For this purpose, we constructed reduced versions of the full MDP described in Section 4.2, where one task factor (such as offer, energy…) was omitted by averaging transition probabilities and rewards across the levels of that factor. For each reduced MDP, we computed the state action value function (*Q_reduced_*) and mapped it back onto the full MDP by assigning the aggregated states’ values to their corresponding original states. Using this mapping, we derived a policy from the reduced MDP and calculated its expected return in the full MDP. This analysis measures the reward sensitivity of each task factor: if ignoring a factor leads to a large loss in return, optimal behavior depends strongly on information about that factor.

Second, we asked whether participants’ preferences themselves define a useful policy in the absence of forward planning. We derived policies from the preference score of the full Preference model and from reduced Preference models in which one task factor was omitted. We then computed the expected return of each preference-based policy relative to the optimal policy. If task-specific preferences provide an adaptive approximation to optimal choice, the full preference policy should outperform a stochastic policy and removing behaviorally important factors should reduce expected return.

Finally, we explored a possible source of these preferences. Within the policy-compression framework, compressed policies can be described in terms of marginal action probabilities (Lai and Gershman 2021). We therefore computed, under the optimal policy, the probability of accepting under each level of each task factor after marginalizing over all other factors. For example, we computed the frequency with which an offer of 1 should be accepted across all energy and cost states. We then compared these factor-specific marginal action probabilities with the fitted preference weights. A close correspondence would suggest that participants’ preferences may approximate feature-specific marginal action tendencies of the optimal policy.

### 4.6 Reaction times analysis

In our task, exact computation of optimal policy is prohibitively expensive, given the size of the task’s state space (1,456 states), the forward planning horizon, and the highly probabilistic nature of the task. Preferences derived from task knowledge or from task experience are presumably readily available to the participants or require minimal computation. A part of our hypothesis is that instead of systematically evaluating all future outcomes before making a choice, participants consider the entropy of their preferences to determine how much they should engage in forward planning. If so, we would expect participants’ RT to reflect the entropy of their preferences, as more certain preferences should be associated with less forward planning, resulting in faster RT.

Ott et al. (2022a) showed that response times were slower when the planning-derived DV for accepting and rejecting were similar, and that this forward planning conflict effect was greater for intermediate than for extreme offers. We reinterpret this context dependence as a consequence of preference entropy. Intermediate offers tend to generate weaker preference signals, whereas extreme offers tend to produce clearer action tendencies. Accordingly, we expected forward planning conflict to slow responses most when preferences were weak.

To test whether the effect of forward planning conflict should depend on preference entropy, we modelled participants’ RTs as a function of forward planning conflict (i.e. *conflict* = −|*DV*|), preference entropy (fitted in the Preference model) and the modulation of forward planning by preference entropy:

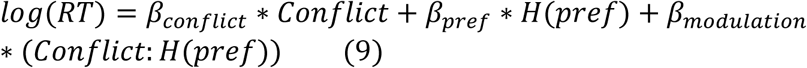

We predicted a positive modulation, such that forward planning conflict would be associated with longer RTs when preference entropy was high, that is, when preferences were weak or uncertain.

## 5 Acknowledgements

The authors thank Qian Chu for their valuable comments and suggestions. This work was funded by the funded by the German Research Foundation (DFG, Deutsche Forschungsgemeinschaft) – 538763959 – and as part of Germany’s Excellence Strategy – EXC 2050/2 – Project ID 390696704 – Cluster of Excellence ‘Centre for Tactile Internet with Human-in-the-Loop’ (CeTI) of TUD Dresden University of Technology.

## 6 Contribution statement

Conceptualization: A.L., F.O., S.K.; Data curation: A.L., F.O.; Formal analysis: A.L.; Funding acquisition: F.O., S.K.; Investigation: A.L.; Methodology: A.L., S.K.; Project management: A.L.; Resources: S.K.; Software: A.L.; Supervision: S.K.; Validation: A.L.; Visualization: A.L., S.K.; Writing - original draft: A.L., S.K.; Writing - review & editing: A.L., F.O., S.K.

## 7 Data and code statement

All data used in this paper were previously published as part of Ott et al. (2022a) and are available at https://zenodo.org/records/6328296. Code can be found on github https://github.com/AlexLepauvre/state_abstraction_paper. In addition, we’ve created a custom toolbox to implement reinforcement learning algorithm, which can be found here: https://github.com/AlexLepauvre/state-abstraction

## 9 Supplementary material

## 9.1 Supplementary figures

**Figure S1:**
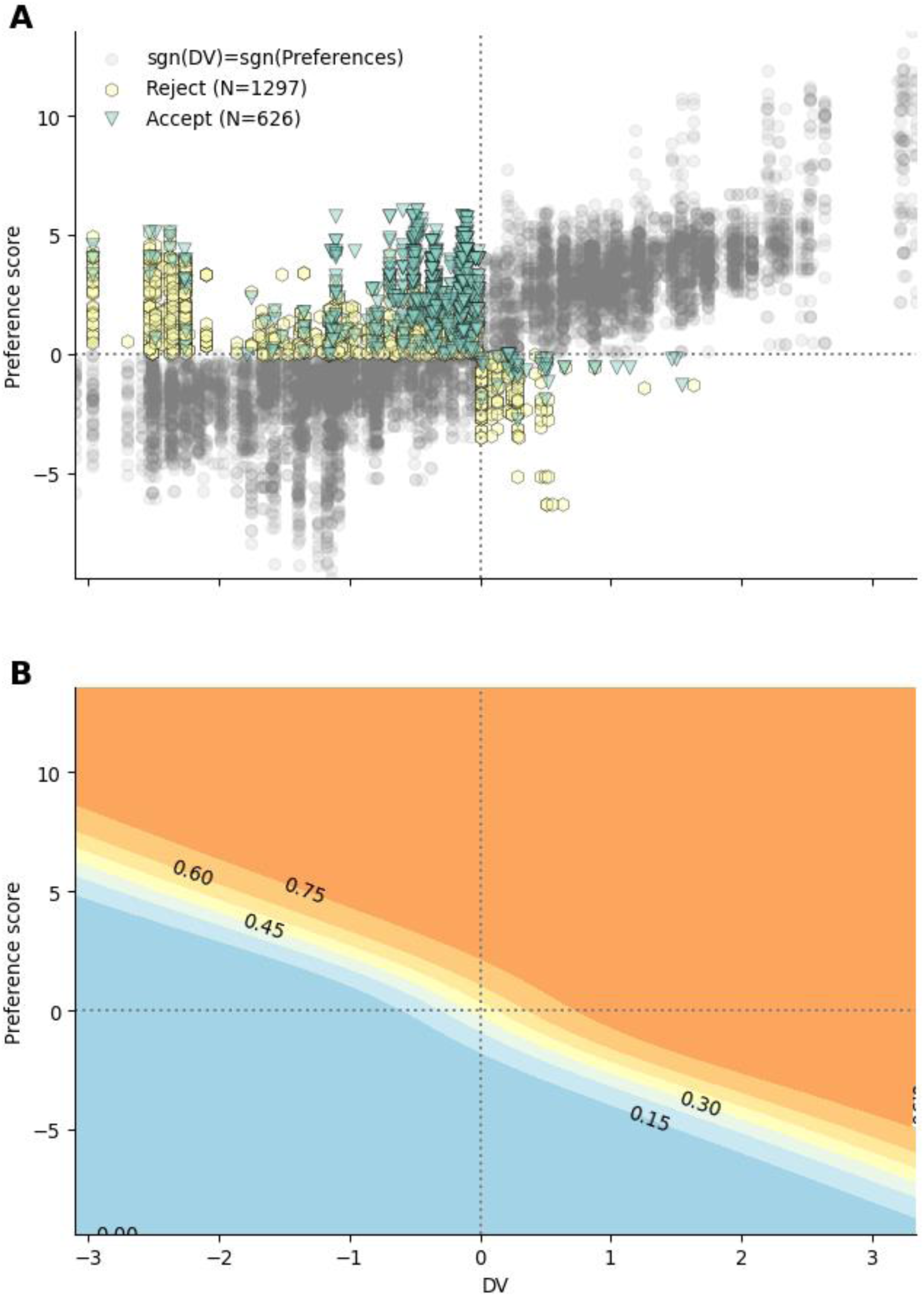
Interaction between preferences and DV. (A) Participants responses as a function of decision values of preference score. The response in the off-diagonal quadrants (top left and bottom right) are color coded per response given, to highlight that participants responses reflect the strongest signal (tend to accept despite DV being negative if preferences are high enough). (B) Fitted probability of accepting as a function of DV and preference score. Contours with annotations represent a probability of accepting. The shape of the contour lines highlight that participants choices reflect the modulation of DV by preference score. The contour trace indicate that when preference and DV are opposed, participants follow the strongest signal.

## 9.2 Supplementary tables

**Table S1.** Main model comparison, comparing the fit of the preference, context, frequency and pure planning models.

|  | elpd_loo | p_loo | elpd_diff | se | dse |
| --- | --- | --- | --- | --- | --- |
| Preference | -1492.47 | 256.484 | 0 | 48.7583 | 0 |
| Context | -1657.96 | 160.738 | 165.486 | 52.6188 | 19.6029 |
| Frequency prior | -2073.86 | 77.4975 | 581.384 | 55.6579 | 35.8832 |
| Planning | -2078.76 | 68.6015 | 586.285 | 55.5599 | 36.001 |

**Table S2.** Summary of the parameters of the preference model. The table displays the mean and 95% highest density interval (HDI) for the population level parameters associated with the planning values, the preference regressors and their interaction.

|  | mean | sd | hdi_2.5% | hdi_97.5% |
| --- | --- | --- | --- | --- |
| $\beta_{plan}$ | 2.082 | 0.248 | 1.603 | 2.577 |
| $\beta_{O=1}$ | -2.371 | 0.987 | -4.194 | -0.381 |
| $\beta_{O=2}$ | -1.611 | 0.955 | -3.537 | 0.17 |
| $\beta_{O=3}$ | 1.656 | 0.958 | -0.092 | 3.611 |
| $\beta_{O=4}$ | 2.955 | 0.985 | 1.161 | 4.979 |
| $\beta_{CC=1}$ | 0.724 | 1.166 | -1.592 | 2.884 |
| $\beta_{CC=2}$ | -0.178 | 1.164 | -2.516 | 1.955 |
| $\beta_{FC=1}$ | 0.416 | 1.165 | -1.866 | 2.711 |
| $\beta_{FC=1}$ | 0.176 | 1.164 | -2.219 | 2.381 |
| $\beta_{E=0}$ | -4.429 | 0.932 | -6.266 | -2.666 |
| $\beta_{E=1}$ | -0.442 | 0.774 | -1.88 | 1.137 |
| $\beta_{E=2}$ | 0.018 | 0.732 | -1.397 | 1.421 |
| $\beta_{E=3}$ | 0.237 | 0.733 | -1.069 | 1.747 |
| $\beta_{E=4}$ | 0.337 | 0.74 | -0.991 | 1.856 |
| $\beta_{E=5}$ | 0.829 | 0.753 | -0.569 | 2.323 |
| $\beta_{E=6}$ | 4.248 | 0.962 | 2.465 | 6.186 |
| $\beta_{interaction}$ | 1.264 | 0.513 | 0.308 | 2.294 |

## 9.3 Supplementary model comparison

### 9.3.1 Preference model comparison

The Preference model was observed to fit the data best, showing that participants behavior reflects the integration of a planning component together with priors reflecting the structure of the task, shedding light on the structure of the priors themselves. To further show that participants priors reflect the whole structure of the task, we modelled the data removing one experimental factor from the preferences fitting one at a time and compared to the full model, to show that our results cannot be explained by the fact that participants’s priors do not reflect a single nor a subset of experimental factors. To further show that participants reliance on planning is modulated by the strength of their preferenecs, a model without the modulatory effect was fitted to highlight the increase in fit associated with the modulatory effect of preferences entropy on DV. Finally, to show that participants response is best explained by a combination of planning and preference component, we also fitted a model with only the preference component omitting the planning regressor.

**Figure S2:**
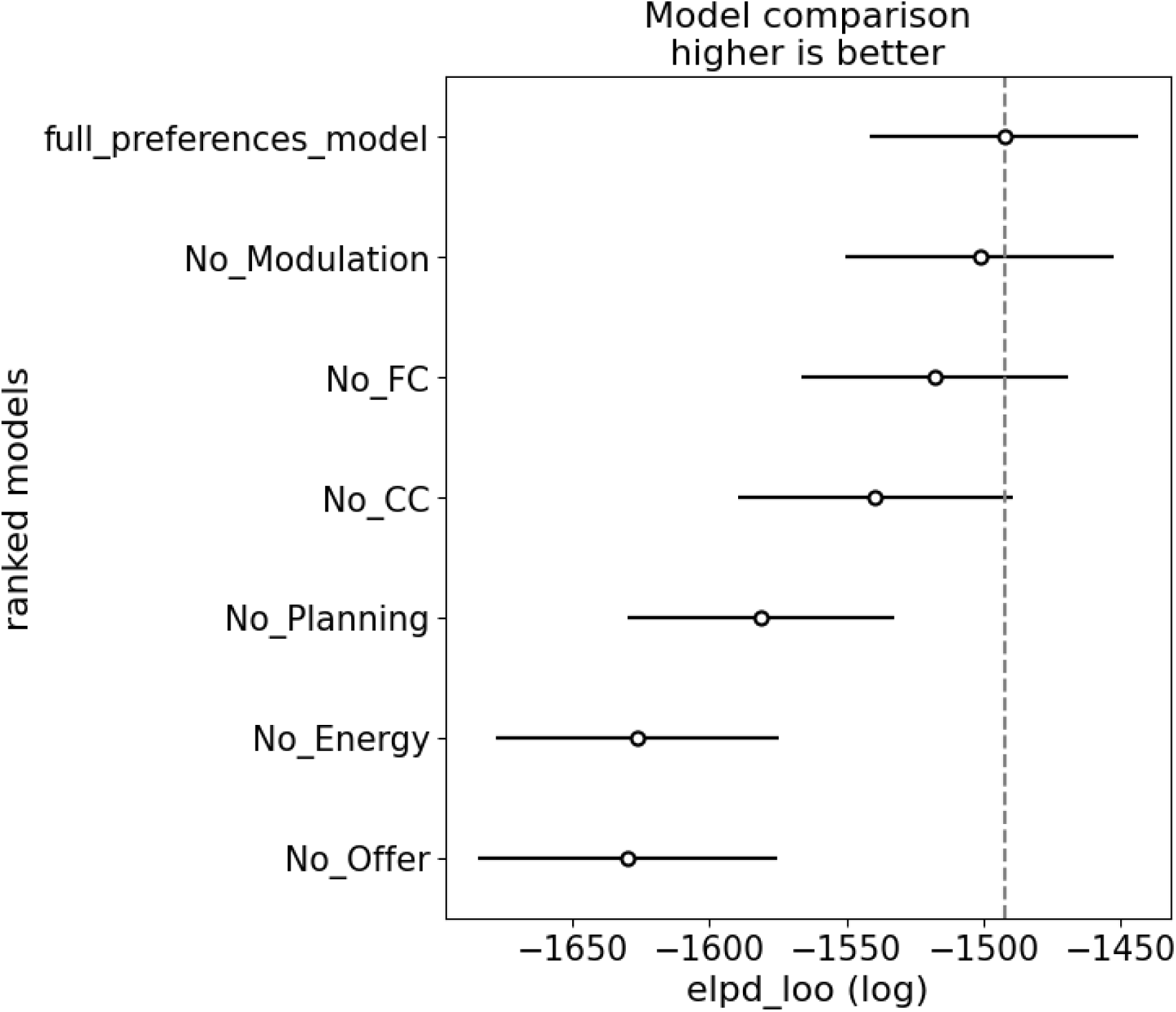
Results of Preference model comparison. Model abbreviations: (No_interaction) model comprising planning and preference component, without interaction between the two, (No_CC) model where current cost factor was omitted from preferences, (No_FC) model where future cost factor was omitted from preferences, (No_Planning) model where DV was omitted, (No_Energy) model where energy factor was omitted from preferences, (No_Offer) model where offer factor was omitted from preferences

**Table S3.** Results of the comparison between different preferences models.

| Results of the comparison between different preferences models |  |  |  |  |  |
| --- | --- | --- | --- | --- | --- |
|  | elpd_loo | p_loo | elpd_diff | se | dse |
| full_preferences_model | -1492.47 | 256.484 | 0 | 48.7583 | 0 |
| No_Modulation | -1501.44 | 245.861 | 8.9671 | 48.9103 | 5.28966 |
| No_FC | -1519.82 | 229.718 | 27.351 | 48.6187 | 7.56828 |
| No_CC | -1539.49 | 262.803 | 47.0195 | 49.8892 | 9.74314 |
| No_Planning | -1581.23 | 217.308 | 88.758 | 48.8039 | 17.3434 |
| No_Energy | -1626.44 | 211.155 | 133.971 | 51.5682 | 17.7893 |
| No_Offer | -1629.84 | 235.203 | 137.365 | 54.4382 | 19.2265 |

### 9.3.2 Frequency model comparison

In the main text, we compared the Preference model with a model in which priors are computed based on action frequency. In this model, the priors are computed using a linearly decaying moving average, discounting actions that occured further in the past. We presented the results obtained when the priors are derived from the 20 previous trials. This number of trials was selected by exploring priors with *k* = [2,5,10,15,20,30,50] to identify the values that fit the data the best:

**Figure S3:**
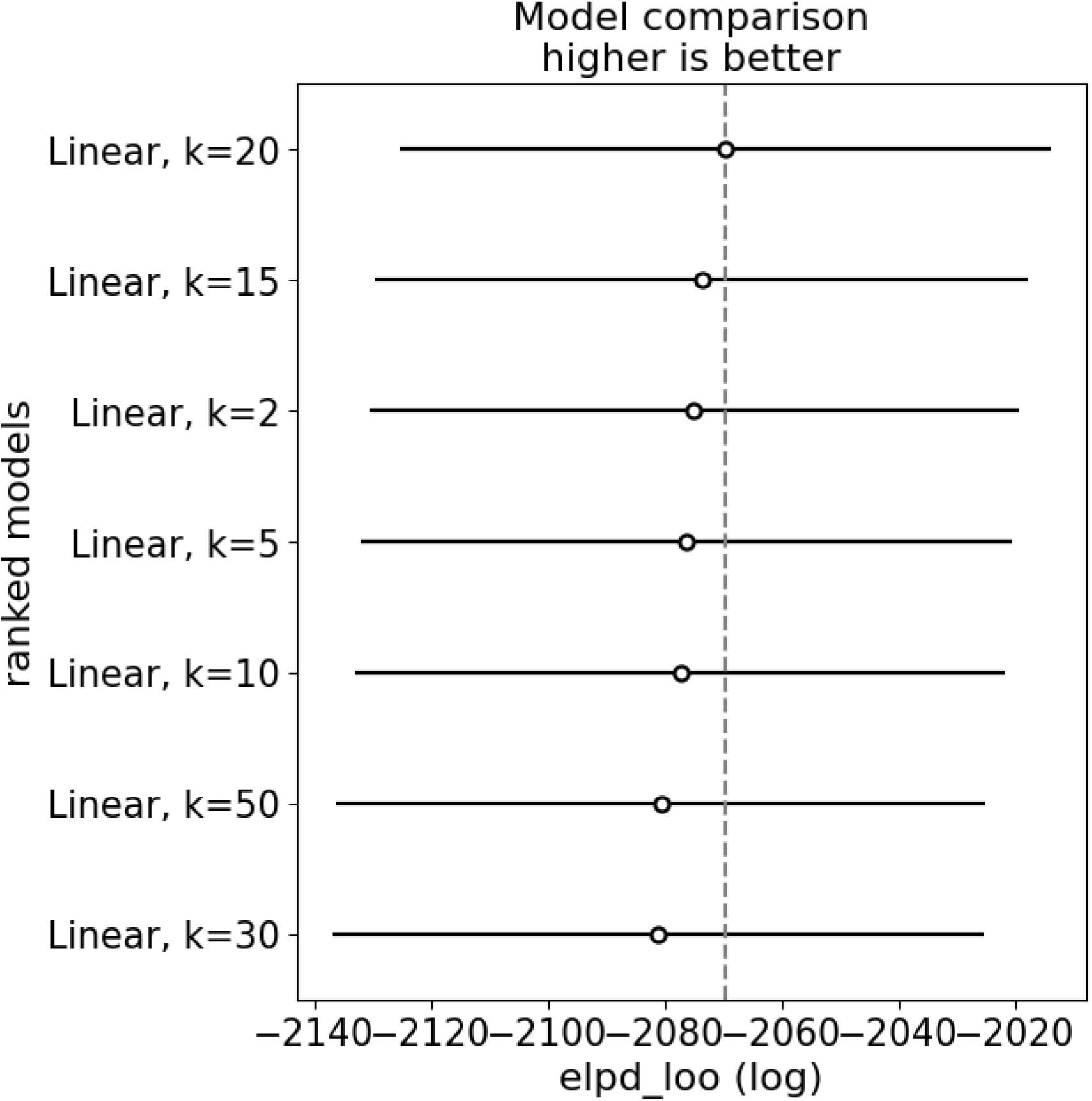
Results of frequency model comparison, comparing the temporal integration window, going back k=[2, 5, 10, 15, 20, 30, 50]

**Table S4.** Results of the frequency model comparisons with different temporal integration windows.

|  | elpd_loo | p_loo | elpd_diff | se | dse |
| --- | --- | --- | --- | --- | --- |
| Linear, k=20 | -2069.6 | 78.3873 | 0 | 55.6858 | 0 |
| Linear, k=15 | -2073.74 | 74.2703 | 4.13799 | 55.8878 | 5.24083 |
| Linear, k=2 | -2074.98 | 75.5677 | 5.37448 | 55.6833 | 6.29665 |
| Linear, k=5 | -2076.3 | 78.84 | 6.69933 | 55.6484 | 5.58139 |
| Linear, k=10 | -2077.34 | 74.0643 | 7.73213 | 55.5341 | 5.65329 |
| Linear, k=50 | -2080.71 | 78.7902 | 11.1052 | 55.7161 | 5.09526 |
| Linear, k=30 | -2081.1 | 75.786 | 11.4922 | 55.7602 | 5.46973 |

### 9.4 Parameters recovery

In the main text, the Preference model was observed to fit participants behavior the best, supporting the hypothesis that participants decision reflects the integration of forward planning with state dependent preferences. The Preference model is rather complex, with preference parameters being estimated as latent variable, and entailing an additional entropy transformation in the interaction term. To establish that despite the complexity of the model, it is still capable of accurately estimate all parameters, we perform a parameter recovery analysis in which we simulate data using the mean parameters recovered from the original data. We then fit the same model to the simulated data. We can then compare the estimated parameters from the simulated data against ground truth to establish the model capacity to identify parameters correctly.

**Figure S4:**
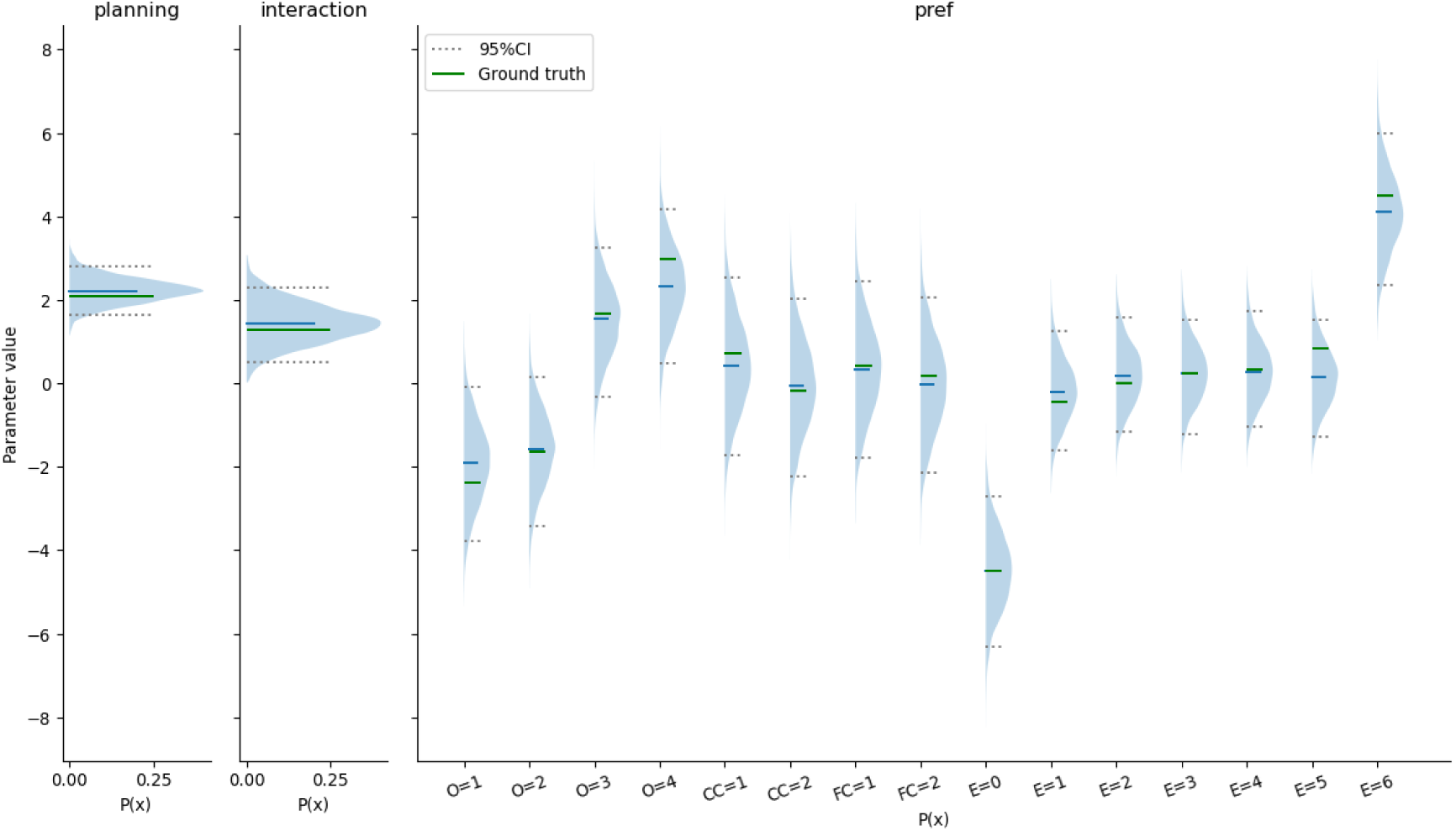
Population level parameters recovery

Across all participants, the ground truth parameters is well within the credible intervals of the retrieved parameters from the data, indicating that our model is indeed capable of recovering the parameters of the underlying generative model.

## Notes

https://zenodo.org/records/6328296

https://github.com/AlexLepauvre/state_abstraction_paper

https://github.com/AlexLepauvre/state-abstraction

### Competing Interest Statement

The authors have declared no competing interest.

## References

Abril-Pla, Oriol, Virgile Andreani, Colin Carroll, et al. 2023. “PyMC: A Modern, and Comprehensive Probabilistic Programming Framework in Python.” PeerJ Computer Science 9: e1516.

Ashburner, John, Gareth Barnes, Chun-Chuan Chen, et al. 2014. “SPM12 Manual.” Wellcome Trust Centre for Neuroimaging, London, UK 2464 (4): 53.

Baayen, R Harald, and Petar Milin. 2010. “Analyzing Reaction Times.” International Journal of Psychological Research 3 (2): 12–28.

Bari, Bilal A, and Samuel J Gershman. 2023. “Undermatching Is a Consequence of Policy Compression.” Journal of Neuroscience 43 (3): 447–57.

Barr, Dale J, Roger Levy, Christoph Scheepers, and Harry J Tily. 2013. “Random Effects Structure for Confirmatory Hypothesis Testing: Keep It Maximal.” Journal of Memory and Language 68 (3): 255–78.

Berthet, Vincent, and Vincent De Gardelle. 2023. “The Heuristics-and-Biases Inventory: An Open-Source Tool to Explore Individual Differences in Rationality.” Frontiers in Psychology 14: 1145246.

Callaway, Frederick, Bas Van Opheusden, Sayan Gul, et al. 2022. “Rational Use of Cognitive Resources in Human Planning.” Nature Human Behaviour 6 (8): 1112–25.

Capretto, Tomás, Camen Piho, Ravin Kumar, Jacob Westfall, Tal Yarkoni, and Osvaldo A Martin. 2022. “Bambi: A Simple Interface for Fitting Bayesian Linear Models in Python.” Journal of Statistical Software 103 (15): 1–29. 10.18637/jss.v103.i15.

Clark, Andy. 2013. “Whatever Next? Predictive Brains, Situated Agents, and the Future of Cognitive Science.” Behavioral and Brain Sciences 36 (3): 181–204.

Friston, Karl. 2010. “The Free-Energy Principle: A Unified Brain Theory?” Nature Reviews Neuroscience 11 (2): 127–38.

Gershman, S. J. (2020). Origin of perseveration in the trade-off between reward and complexity. Cognition, 204, 104394.

Gershman, Samuel J, Eric J Horvitz, and Joshua B Tenenbaum. 2015. “Computational Rationality: A Converging Paradigm for Intelligence in Brains, Minds, and Machines.” Science 349 (6245): 273–78.

Gershman, Samuel J, and Armin Lak. 2025. “Policy Complexity Suppresses Dopamine Responses.” Journal of Neuroscience 45 (9).

Gigerenzer, Gerd, and Henry Brighton. 2009. “Homo Heuristicus: Why Biased Minds Make Better Inferences.” Topics in Cognitive Science 1 (1): 107–43.

Gigerenzer, Gerd, Peter M Todd, the ABC Research Group, et al. 2000. Simple Heuristics That Make Us Smart. Oxford University Press.

Gilovich, Thomas, and Dale W Griffin. 2002. Heuristics and Biases: The Psychology of Intuitive Judgment. Cambridge university press.

Gluth, Sebastian, Jörg Rieskamp, and Christian Büchel. 2012. “Deciding When to Decide: Time-Variant Sequential Sampling Models Explain the Emergence of Value-Based Decisions in the Human Brain.” Journal of Neuroscience 32 (31): 10686–98.

Ho, Mark K, David Abel, Carlos G Correa, Michael L Littman, Jonathan D Cohen, and Thomas L Griffiths. 2022. “People Construct Simplified Mental Representations to Plan.” Nature 606 (7912): 129–36.

Huys, Quentin JM, Neir Eshel, Elizabeth O’Nions, Luke Sheridan, Peter Dayan, and Jonathan P Roiser. 2012. “Bonsai Trees in Your Head: How the Pavlovian System Sculpts Goal-Directed Choices by Pruning Decision Trees.” PLoS Computational Biology 8 (3): e1002410.

Knill, David C, and Alexandre Pouget. 2004. “The Bayesian Brain: The Role of Uncertainty in Neural Coding and Computation.” TRENDS in Neurosciences 27 (12): 712–19.

Körding, Konrad P, and Daniel M Wolpert. 2004. “Bayesian Integration in Sensorimotor Learning.” Nature 427 (6971): 244–47.

Kuperwajs, Ionatan, and Wei Ji Ma. 2021. “Planning to Plan: A Bayesian Model for Optimizing the Depth of Decision Tree Search.” Proceedings of the Annual Meeting of the Cognitive Science Society 43.

Lai, Lucy, and Samuel J Gershman. 2021. “Policy Compression: An Information Bottleneck in Action Selection.” In Psychology of Learning and Motivation, vol. 74. Elsevier.

Lai, Lucy, and Samuel J Gershman. 2024. “Human Decision Making Balances Reward Maximization and Policy Compression.” PLOS Computational Biology 20 (4): e1012057.

Lai, Lucy, Ann Zixiang Huang, and Samuel J Gershman. 2022. “Action Chunking as Policy Compression.” PsyArXiv.

Lee, Douglas G, and Jean Daunizeau. 2021. “Trading Mental Effort for Confidence in the Metacognitive Control of Value-Based Decision-Making.” Elife 10: e63282.

Lee, Tai Sing, and David Mumford. 2003. “Hierarchical Bayesian Inference in the Visual Cortex.” Journal of the Optical Society of America A 20 (7): 1434–48.

Liu, Shuze, and Samuel Joseph Gershman. 2025. “Action Subsampling Supports Policy Compression in Large Action Spaces.” PLOS Computational Biology 21 (9): e1013444.

Liu, S., Kikumoto, A., Badre, D., & Gershman, S. J. (2025). Neural and behavioral signatures of policy compression in cognitive control. Cerebral Cortex, 35(8), bhaf223.

Ott, Florian, Eric Legler, and Stefan J Kiebel. 2022a. “Forward Planning Driven by Context-Dependant Conflict Processing in Anterior Cingulate Cortex.” NeuroImage 256: 119222. 10.1016/j.neuroimage.2022.119222.

Ott, Florian, Eric Legler, and Stefan J Kiebel. 2022b. Forward Planning Driven by Context-Dependent Conflict Processing in Anterior Cingulate Cortex - Analysis Code and Datasets. V. 1.2. Zenodo, released. 10.5281/zenodo.6328296.

Ott, Florian, Dimitrije Marković, Alexander Strobel, and Stefan J Kiebel. 2020. “Dynamic Integration of Forward Planning and Heuristic Preferences During Multiple Goal Pursuit.” PLoS Computational Biology 16 (2): e1007685.

Parush, Naama, Naftali Tishby, and Hagai Bergman. 2011. “Dopaminergic Balance Between Reward Maximization and Policy Complexity.” Frontiers in Systems Neuroscience 5: 22.

Pouget, Alexandre, Jeffrey M Beck, Wei Ji Ma, and Peter E Latham. 2013. “Probabilistic Brains: Knowns and Unknowns.” Nature Neuroscience 16 (9): 1170–78.

Rahnev, Dobromir, and Rachel N Denison. 2018. “Suboptimality in Perceptual Decision Making.” Behavioral and Brain Sciences 41: e223.

Ruge, Hannes, and Uta Wolfensteller. 2010. “Rapid Formation of Pragmatic Rule Representations in the Human Brain During Instruction-Based Learning.” Cerebral Cortex 20 (7): 1656–67.

Schwartenbeck, Philipp, Thomas HB FitzGerald, Christoph Mathys, Ray Dolan, and Karl Friston. 2015. “The Dopaminergic Midbrain Encodes the Expected Certainty about Desired Outcomes.” Cerebral Cortex 25 (10): 3434–45.

Schwöbel, Sarah, Dimitrije Marković, Michael N Smolka, and Stefan Kiebel. 2024. “Joint Modeling of Choices and Reaction Times Based on Bayesian Contextual Behavioral Control.” PLoS Computational Biology 20 (7): e1012228.

Schwöbel, Sarah, Dimitrije Marković, Michael N Smolka, and Stefan J Kiebel. 2021. “Balancing Control: A Bayesian Interpretation of Habitual and Goal-Directed Behavior.” Journal of Mathematical Psychology 100: 102472.

Silver, David, Aja Huang, Chris J Maddison, et al. 2016. “Mastering the Game of Go with Deep Neural Networks and Tree Search.” Nature 529 (7587): 484–89.

Singmann, Henrik, and David Kellen. 2019. “An Introduction to Mixed Models for Experimental Psychology.” In New Methods in Cognitive Psychology. Routledge.

Sutton, Richard S, Andrew G Barto, et al. 1998. Reinforcement Learning: An Introduction. Vol. 1. MIT press Cambridge.

Tishby, Naftali, and Daniel Polani. 2010. “Information Theory of Decisions and Actions.” In *Perception-Action Cycle: Models*, Architectures, and Hardware. Springer.

Tsetsos, Konstantinos, Rani Moran, James Moreland, Nick Chater, Marius Usher, and Christopher Summerfield. 2016. “Economic Irrationality Is Optimal During Noisy Decision Making.” Proceedings of the National Academy of Sciences 113 (11): 3102–7.

Tversky, Amos, and Daniel Kahneman. 1974. “Judgment Under Uncertainty: Heuristics and Biases: Biases in Judgments Reveal Some Heuristics of Thinking Under Uncertainty.” Science 185 (4157): 1124–31.

Vehtari, Aki, Andrew Gelman, and Jonah Gabry. 2017. “Practical Bayesian Model Evaluation Using Leave-One-Out Cross-Validation and WAIC.” Statistics and Computing 27 (5): 1413–32.

